# Development of Synthetic Modulator Enabling Long-Term Propagation and Neurogenesis of Human-Derived Neural Progenitor Cells

**DOI:** 10.1101/2023.05.15.540695

**Authors:** Ceheng Liao, Ying Guan, Jihui Zheng, Xue Wang, Meixia Wang, Zhouhai Zhu, Qiyuan Peng, Hong-Hui Wang, Meng Li

## Abstract

Neural progenitor cells (NPCs) are important cells for in vitro drug screening and the cell-based therapy for brain-related disorders, which requires well-defined and reproducible culture systems. Current strategy the use of protein growth factors presents challenges in terms of reproducibility and cost. In this study, we have developed a novel DNA-based modulator to regulate FGFR signaling of NPCs, enabling maintenance of the stemness over 50 passages and neurogenesis towards neurons. The DNA-based FGFR-agonist effectively promoted FGFR1 phosphorylation and activated the downstream ERK signaling pathway in FGFR1-positive cells. Using human embryonic stem cell lines, we differentiated them into NPCs and replaced basic fibroblast growth factor (bFGF) in the regulator culture medium with DNA-based FGFR-agonist for artificially elicited FGFR signaling. The results demonstrated that the FGFR-agonist could promote NPCs proliferation and neurosphere formation, recapitulating the function of bFGF. Notably, transcriptomic analysis revealed that FGFR-agonist could customize the stemness-associated transcription program, while decouples the neuronal differentiation program, highly resembling that the native ligand, bFGF. Moreover, our culture condition facilitated the successful propagation of NPCs for over 50 passages, while retaining their ability to efficiently differentiate into neurons. Overall, our approach provides a highly effective method for expanding NPCs, offering new opportunities for disease-in-dish research and drug screening for neural degeneration.

## Introduction

Neurological disorders, such as Alzheimer disease, Parkinson disease, and traumatic brain injuries, pose significant global health challenges, affecting millions of individuals worldwide and leading to substantial impairment in the quality of life for both patients and their families [1-3]. Understanding the underlying pathology of these disorders and developing effective therapies require a comprehensive understanding of neural cell behavior and the ability to manipulate and study neural progenitor cells (NPCs) [4, 5]. NPCs are a crucial cell population in the field of neuroscience research and regenerative medicine due to their unique properties, including self-renewal and the ability to differentiate into various cell types within the nervous system, including neurons, astrocytes, and oligodendrocytes [6, 7]. This remarkable versatility makes NPCs invaluable for a wide range of applications, such as studying disease mechanisms, conducting drug screenings, and developing cell-based therapies to treat or even reverse the effects of neurological disorders[8-10] .

The practical application of NPCs in disease modeling and regenerative medicine relies heavily on the ability to culture and expand these cells on a large scale. Establishing reliable and scalable culture systems for NPCs is essential to ensure a sufficient supply of cells for experimentation and therapeutic purposes. Large-scale culturing of NPCs offers the potential to generate high-quality cellular models that accurately recapitulate disease-associated features, providing valuable insights into disease mechanisms and enabling the development of novel therapeutic interventions [11, 12]. Moreover, the ability to culture and expand NPCs on a large scale allows scientists to conduct extensive drug screenings, testing the efficacy and safety of potential treatments in a controlled and reproducible manner[13, 14] . By establishing reliable and scalable culture systems for NPCs, researchers can unlock the full potential of these versatile cells, paving the way for groundbreaking discoveries and therapeutic advancements in the field of neuroscience and regenerative medicine[15] .

The expansion and maintenance of NPCs in the field of neuroscience research and regenerative medicine face significant challenges, such as establishing a well-defined, reproducible culture system and managing high production costs associated with traditional methods [16, 17]. Conventional NPC culture systems often rely on serum-containing media and protein growth factors like fibroblast growth factor (FGF), epidermal growth factor (EGF), and brain-derived neurotrophic factor (BDNF), which play a vital role in sustaining NPCs’ self-renewal and differentiation capacities [18, 19]. However, these methods present challenges, such as batch-to-batch variability in serum-containing media [20], potential immunogenicity [21], and elevated production costs [22]. To address these challenges, researchers are exploring alternative approaches, such as chemically defined media and small molecules as substitutes for growth factors, aiming to establish a more consistent and cost-effective culture system while preserving NPCs’ stem cell properties and differentiation potential [23, 24]. These promising strategies, such as the development of chemically defined media and synthetic substitutes for serum or growth factors, offer improved consistency, reproducibility, and cost-effectiveness by providing greater control over the culture environment, eliminating batch-to-batch variability, and reducing the risk of immunogenic responses[25, 26]. Therefore, the development of chemically defined media would facilitate the establishment of NPCs culture would ultimately improve the reproducibility and increasing versatility for exploring their applications in disease modeling and therapeutics, however, remains largely unavailable.

In this study, we developed a novel approach that utilizes a DNA-based modulator to maintain the stemness of NPCs and facilitate the expansion of NPCs (**Scheme 1**). By replacing protein-based native growth factors with DNA-based modulators, we aimed to overcome the limitations of current NPCs culture systems, such as experimental variability and high production costs[27, 28]. We investigated the potential of artificial DNA-based modulator as a viable alternative to bFGF to maintain and expand human-derived NPCs and decouple the default neuronal differentiation program. To achieve this, we conduct a comprehensive evaluation of DNA modulator mediated FGFR signaling and transcriptome analysis, and assess the maintenance of stemness in NPCs. By utilizing FGFR-agonist, we established a chemically defined culture system that effectively preserves the stem cell characteristics of NPCs. We expect this novel approach has the potential to revolutionize the field of neuroscience research and regenerative medicine by offering a more consistent, scalable, and cost-effective method for the large-scale production of NPCs.

**Scheme 1.**
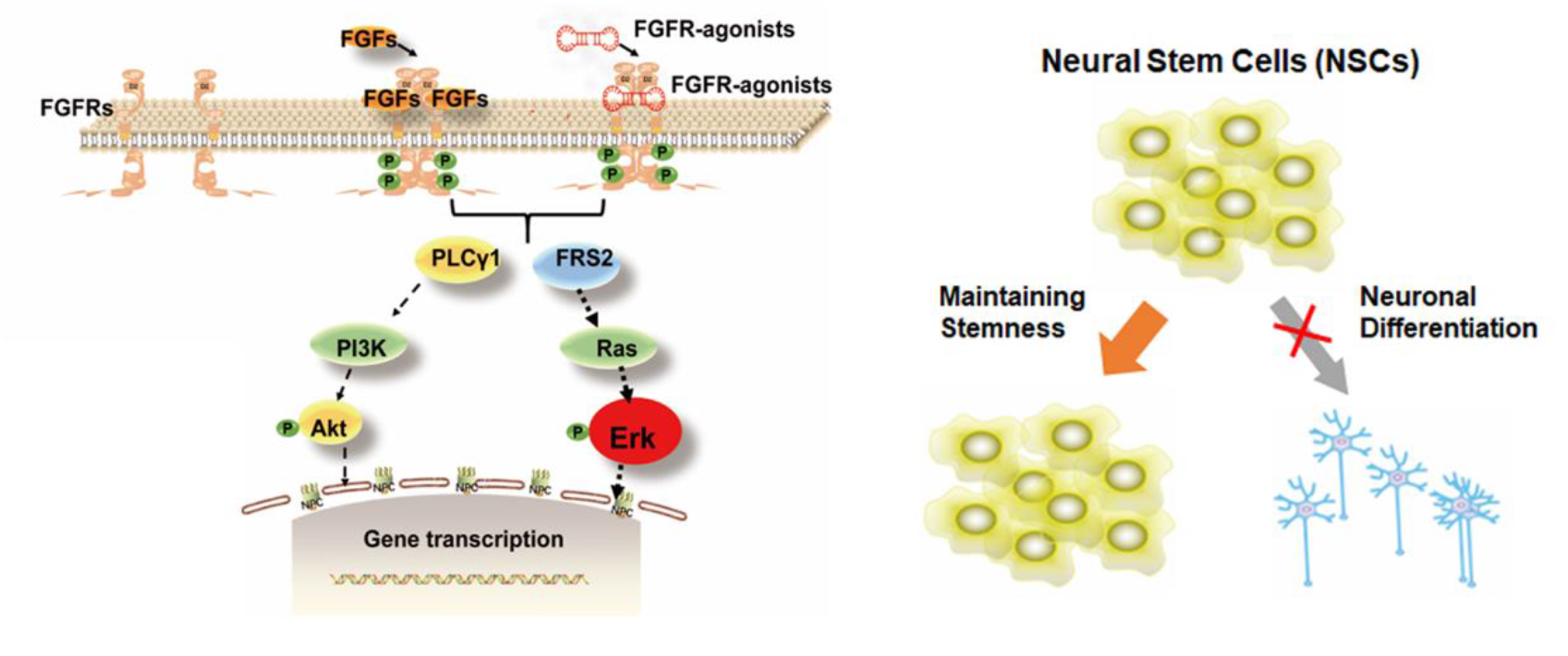
DNA-based FGFR-agonist for NPCs maintenance

## Results

### Design and characterization of DNA-based FGFR-agonist

We developed a DNA-based synthetic agonist to mimic the function of bFGF, allows for receptor dimerization in a controllable manner to effectively activate the receptor. In this work, an artificial agonist was rationally designed to bind and activate FGF receptor 1 (FGFR1). The essential role of FGF-FGFR1 signaling in NPCs have been demonstrated and emerged development of the bivalent FGFR1-agonist for maintain and support the pluripotency of human embryonic stem cells[29, 30]. Similar strategy to develop DNA-based agonists have been demonstrated to mimic hepatic growth factor (HGF) and vascular endothelia growth factor (VEGF)[31, 32]. In current study, we asked if the bivalent DNA architecture containing two FGFR1-binder could elicit FGFR1 dimerization for signaling propagation in NPCs. We chose a previously characterized monomeric FGFR-binding aptamer as the FGFR-binder, which is a 38-mer stem-loop oligonucleotide (**Fig. 1A, Table 1**). To achieve ligand-induced receptor dimerization, we conjugated two monomeric FGFR1-binder to construct bivalent ligand, which can assembly two cell surface FGFR1 as an FGFR1 agonist. Based on prediction, the bivalent construction of both FGFR1-binder did not affect the secondary structure of the each FGFR1-binding domain (**Fig. 1A**). Using 8% Native PAGE gel electrophoresis, we showed that the molecular weight of single-stranded FGFR-binder was around 25 nt, while the molecular weights of both the FGFR-agonist and its control oligo were approximately 72 nt, probably due to the formed secondary structure (**Fig. 1B**). In comparison, the control oligo with scramble sequence of FGFR-agonist exhibited higher molecular weight, suggesting the diminished structure that is essential for FGFR1 interaction.

**Fig. 1:**
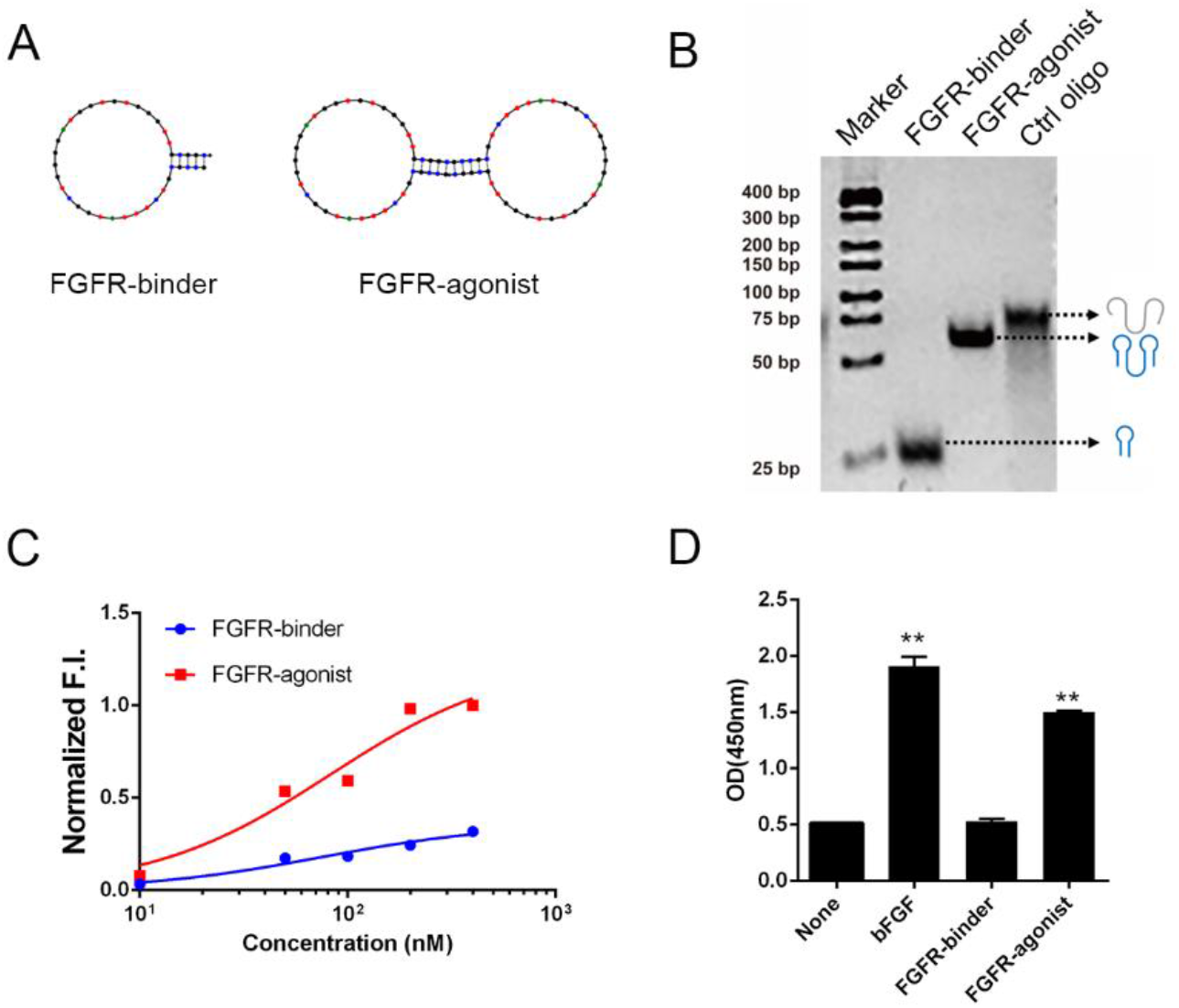
Design and characterization of the FGFR-Agonist. A) The secondary structure of the FGFR-agonist is shown, determined using Mfold software, which indicates the formation of a stem-loop structure. B) The secondary structure of the DNA-based FGFR-agonist is verified using native polyacrylamide gel electrophoresis (PAGE). This panel also includes the determined molecular weights of the single-stranded FGFR-binder, FGFR1-agonist, and a control oligonucleotide (Ctrl oligo). C) The relative binding affinities of the FGFR-binder and FGFR-agonist to FGFR-positive NIH3T3 cells, as determined through flow cytometry (refer to **Fig. S1** for additional details). D) An evaluation of FGFR phosphorylation in NIH3T3 cells in response to the FGFR-agonist, assessed using ELISA. All data are presented as means ± standard deviation (SD) from three independent experiments. **P < 0.0001 (unpaired two-tailed Student’s t-test).

We further carried out flow cytometry analysis to evaluate the cell binding activity of FGFR-agonist using a FGFR1-positive NIH3T3 cell line, the murine fibroblast cell line. The result demonstrated that the FGFR1-binder exhibited high affinity for FGFR1 with a dissociation content of 39 nM (**Fig. 1C**). Notably, the bivalent FGFR1-binder could affect the cell-binding affinity comparing with the monomeric FGFR-binder, while significantly increasing the binding fraction. This data indicate that bivalent construction increases the avidity of the agonist which would be highly useful for FGFR activation. We asked if the FGFR-agonist could activate FGFR phosphorylation using specific antibody against phosphorylated FGFR1 at tyrosine 734 in the western blotting experiment. The result showed that the FGFR-agonist significantly promote the phosphorylation level of FGFR1 compared with the controls treated with monomeric FGFR-binder (**Fig. 1D** and **Fig. S1**). The stability of the FGFR-agonist under physiological conditions in vivo would be highly important for robust effect for in vivo application. To characterize this feature, we carried out a serum stability assay by incubating the FGFR-agonist with 10% FBS at 37°C. After incubation for different duration, we analyzed all samples using 8% Native PAGE gel electrophoresis. The results showed that the FGFR-agonist remained stable after 16 hours in 10% FBS, indicating its suitability for use in subsequent experiments (**Fig. S2**). Together, we successfully constructed a serum-stable DNA-based FGFR-agonist with high binding affinity for FGFR1 and efficient activation of FGFR phosphorylation. This finding paves the way for further exploration of the FGFR-agonist potential in modulating FGF-FGFR signaling pathway.

To explore the role of FGFR-agonist in activating the FGF-FGFR signaling and cellular functions, we chose FGFR-positive ATDC5 cell, which is a murine chondrocyte line that can be activated by bFGF [33]. The serum-starved ATDC5 cells were stimulated with FGFR-agonist (40 nM) and bFGF (20 ng/mL) for 15 min and the phosphorylation of FGFR1 (tyrosine 673) were evaluated using Western blotting. Similar to the function of bFGF, FGFR-agonist significantly increased FGFR1 phosphorylation and ERK1/2 phosphorylation compared to the untreated cells or ctrl oligo-treated cells (**Fig. 2A**). Furthermore, we studied the duration of downstream p-ERK signals induced by FGFR-agonist. The p-ERK signals were detected as early as 5 minutes, decreased by half at 10 minutes, and remained detectable for 60 minutes, showing a decay thereafter (**Fig. 2B** and **Fig. S3**). Interestingly, bFGF induced a more sustained ERK phosphorylation than FGFR-agonist, probably due to the various FGFR1 endocytosis alter the kinetics of downstream signaling [34]. Inspired by the demonstrated role of FGF/FGFR/ERK signal axis to regulate essential behaviors, we next asked if FGFR-agonist affects cell proliferation and the result revealed that FGFR-agonist promoted the cell proliferation comparable to that induced by bFGF (**Fig. 2B**).

**Fig. 2:**
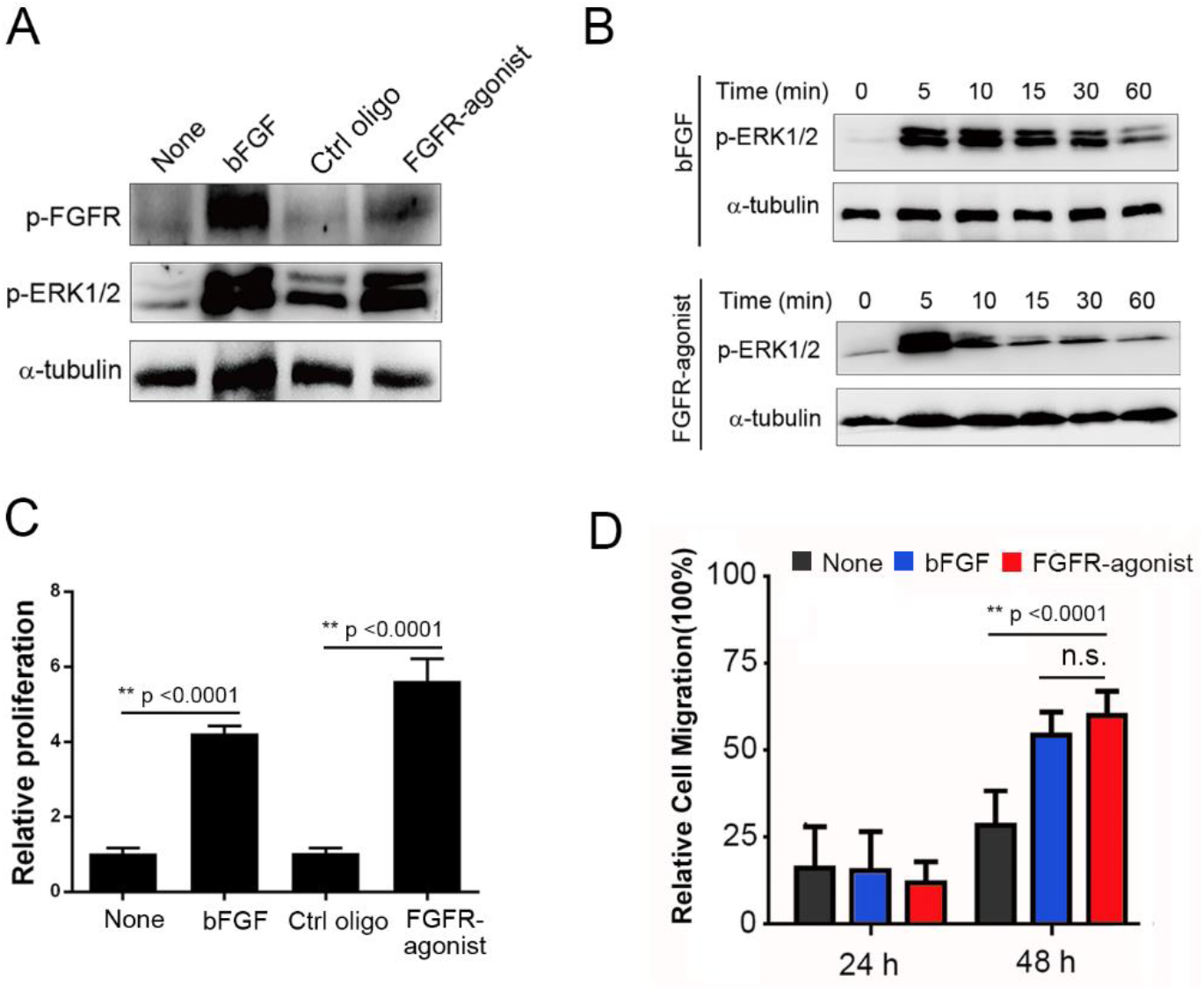
The Efficacy of FGFR-Agonist versus bFGF in FGFR Signaling and Cell Proliferation. A) Assessment of the p-FGFR1(Tyr653/654) and p-ERK1/2 (Thr202/Tyr204) levels in ATDC5 cells through Western blotting, following treatment with either bFGF or the FGFR-agonist. B) Analysis of the time-dependent activation of p-ERK1/2 in ATDC5 cells after exposure to bFGF (20 ng/mL) or the FGFR-agonist (40 nM) over different time intervals (5, 10, 15, 30, 60 minutes). C) A study of ATDC5 cell proliferation under various conditions (None, bFGF, Ctrl oligo, FGFR-agonist), with continuous culture for 2 days. Cell proliferation was measured using the Cell Counting Kit-8 (CCK8). Data are presented as mean ± SD (n = 5), with **P < 0.0001 (based on an unpaired two-tailed Student’s t-test). D) Examination of the impact of the FGFR-agonist on ATDC5 cell migration. Cells were treated with bFGF, the FGFR-agonist, or the FGFR-binder for 2 days. Wound-closure events were tracked using light microscopy, with images captured at 0 h, 24 h, and 48 h. The relative wound closure rates under different experimental conditions were quantified and normalized to establish relative cell migration in comparison with the areas at the start points. Data are presented as means ± SD (n = 3), wit,h **P < 0.0001 and n.s. indicating no significance (based on an unpaired two-tailed Student’s t-test).

We further investigated the effect of the FGFR-agonist on the migratory behavior using a wound scratch assay. The artificial wounds were made by scratching the confluent ATDC5 cells after serum starvation, and then captured images at 0 h, 24 h, and 48 h under different treatment conditions (**Fig. S4**). We found that the cells stimulated with FGFR-agonist achieved a wound closure rate of approximately 40% at 24 hours and around 60% at 48 hours, which was significantly higher than the untreated cells (**Fig. 2C**). Nevertheless, FGFR-agonist exhibited similar activity to endow cells with enhanced migration as observed with bFGF stimulation. Together, we have established the role of FGFR-agonist in modulate FGFR/ERK signaling independent of native bFGF ligand, which could promote proliferative and migratory behaviors of the FGF-responsive cells, which would be highly applicable for the development of innovative culture system to maintain NPCs.

### FGFR-agonist recapitulates the function of bFGF for NPCs maintenance

We next obtained human-derived NPCs according to a previous described pipeline with different combinations of culturing medium (**Fig. S5**). Immunofluorescence analysis confirmed that these cells were positively stained with Nestin and Pax 6, both were NPC markers (**Fig. S6**). Upon neuronal differentiation without bFGF, NPCs could be further differentiated into MAP2-positive neurons (**Fig.S7**). The NPCs were then cultured in dishes without collagen coating and placed the dishes on a shaker to ensure that the cells remained suspended. The suspension growth of NPCs by comparing the growth with and without the addition of bFGF (20 ng/mL) or FGFR-agonist with two different concentrations (40 nM or 200 nM). The bFGF-maintained NPCs exhibited classic characteristic morphology of neurosphere formation from isolated NPCs during 10-day culturing (**Fig. 3A**). In contrast, the untreated group formed cell clusters from single cells within the first 5 days, but the growth trend of the cell clusters dramatically lagged behind the bFGF group. Quantitative results showed that the size of cell clusters in the FGFR-agonist group was similar to that of the bFGF-treated (**Fig. 3B**). After 5 days, the growth trend of neurosphere in the bFGF and FGFR-agonist (40 nM) significantly differed from the untreated group. Until day 10, the volume of neurosphere continued to grow, while the neurosphere in the blank control group dispersed. Higher concentration of FGFR-agonist Our finding suggests that the FGFR-agonist can mimic the function of bFGF, maintaining the neurosphere growth.

**Fig. 3:**
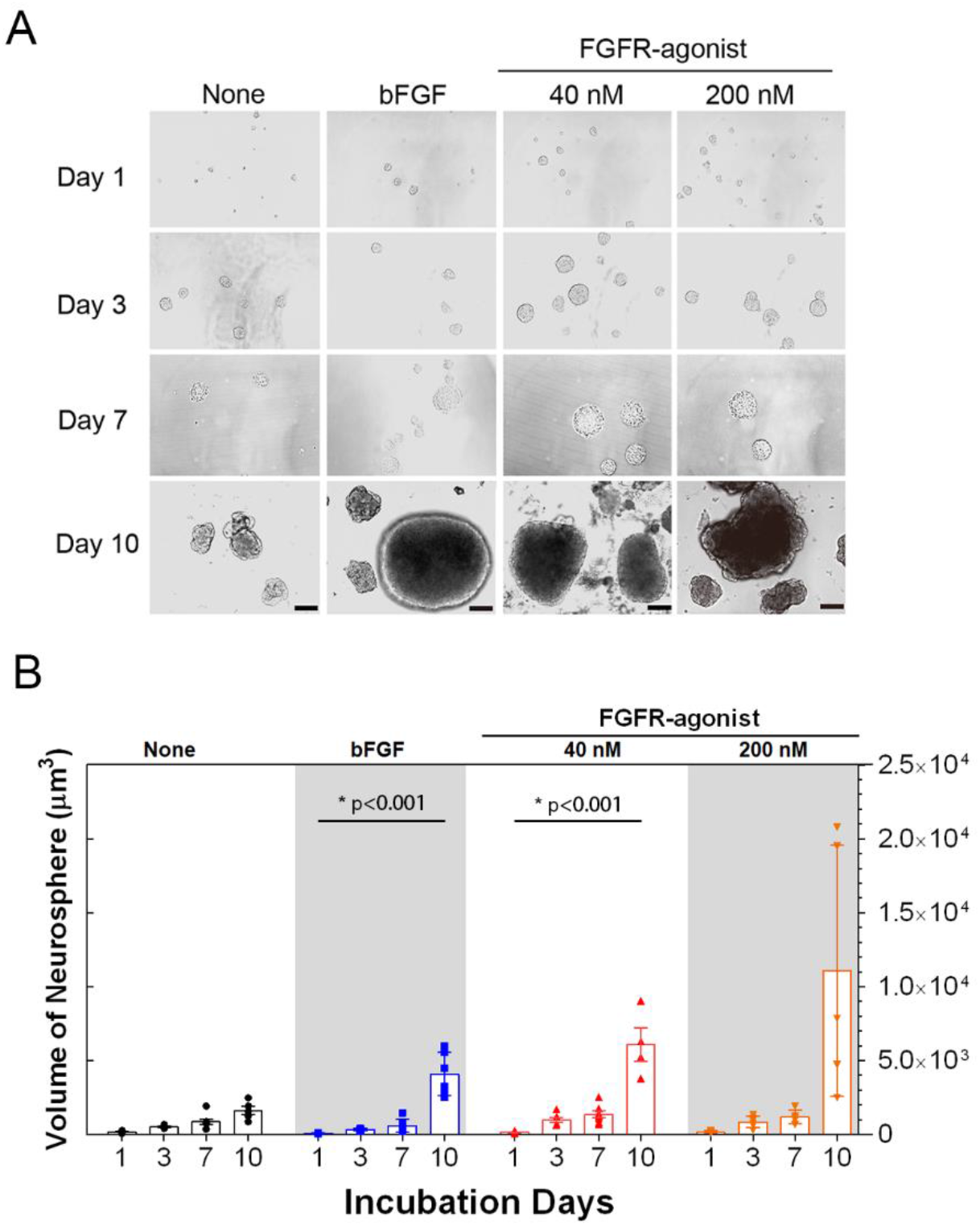
FGFR-Agonist Preserves Self-Renewal and Stemness in Cultured NPCs. (A) Showcases representative images of neurospheres generated by NPCs, which were cultured in suspension for 10 days with or without the presence of bFGF (20 ng/mL), FGFR-agonist (40 nM)or FGFR-agonist (200 nM). The scale bar represents 100 μm. (B) This panel presents a quantification of the changes in neurosphere sizes over time, with or without indicated treatments. Data are displayed as mean ± SD (n = 3), with *P < 0.001 (based on a two-way ANOVA).

### FGFR-agonist affect the transcriptome program of NPCs

To further investigate the effects of the FGFR-agonist in maintaining NPCs, we performed nanopore sequencing and bioinformatic analysis on the transcriptome of NPCs treated with the FGFR-agonist and bFGF. Total RNA from neurosphere maintained in FGFR-agonist and bFGF for 10 days were extracted and then subjected to nanopore sequencing, which offers long-read sequencing capabilities and the ability to detect full-length transcripts. The obtained sequencing data were subsequently processed and analyzed bioinformatically to identify differentially expressed genes (DEGs) and investigate their functional significance. A total of 111 genes were found to be upregulated, while 97 genes were downregulated in NPCs treated with the FGFR-agonist compared to the untreated group (**Table S2**). The volcano plot of differentially expressed genes (DEGs) clearly demonstrated the significant differential expression of transcripts between the FGFR-agonist-treated group and the untreated group (**Fig. 4A**). Functional enrichment analysis of the upregulated DEGs showed significant enrichment in gene ontology (GO) terms related to stem cell maintenance, including “signaling pathways regulating pluripotency of stem cells”, “MAPK signaling pathway”, “Wnt signaling pathway”, “PI3K-Akt signaling pathway” and “Hippo signaling pathway” (**Fig. 4B**). Particularly, the upregulated DEGs revealed significant enrichment in signaling pathways associated with NPC maintenance and self-renewal, including the “Forebrain development” “Regulation of ERK1 and ERK2 cascade” and “Cellular responses to platelet-derived growth factor stimulus” (**Fig. 4C**). Particularly, we focused on the KEGG pathway of regulating pluripotency of stem cells (ko04550) and the analysis highlighted the upregulation of several pivotal neural stem cell markers in the FGFR-agonist treated group (**Fig. S8**). Several key genes were identified to display upregulated expression following the FGFR-agonist treatment, including FGFR, MERK, ERK1/2, Jarid2, SOX2, and c-Myc. Notably, SOX2 and c-Myc, are both instrumental transcription factors in promoting self-renewal and pluripotency of embryonic or neural stem cells[35, 36]. Interestingly, the KEGG pathway enrichment analysis of the downregulated genes upon FGFR-agonist treatment showed that the top five related neuronal diseases were associated with the neuronal dysfunctions, such as Alzheimer disease, Parkinson disease, Huntington’s disease, and Amyotrophic lateral sclerosis, validating the potentials of FGFR-agonist to decouple the mature neuronal function (**Fig. 4D**). These findings underline the potential of FGFR-agonist in promoting and preserving the stemness of neural progenitor cells.

**Fig. 4:**
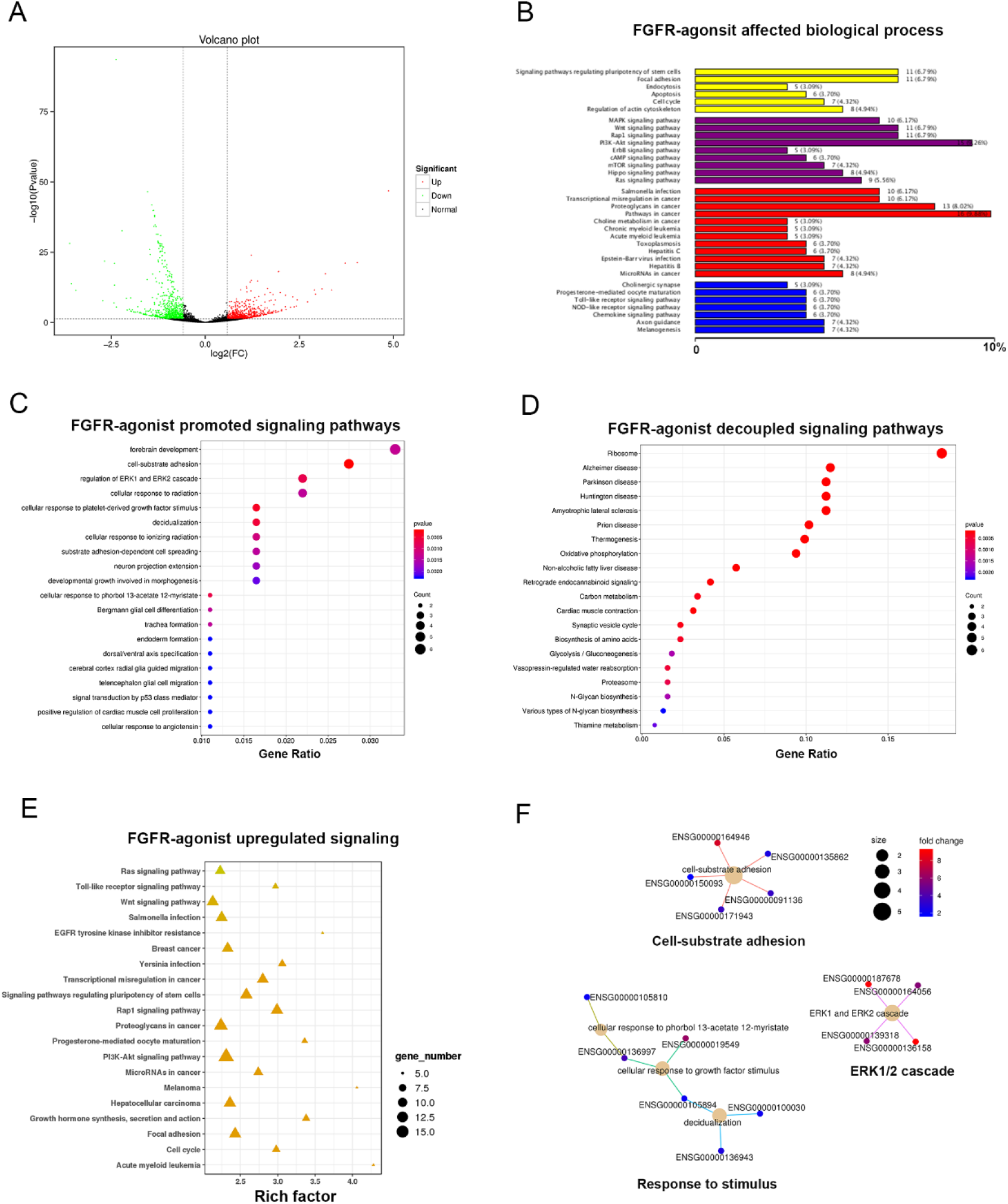
FGFR-agonist programmed transcriptional profile of the NPCs. (A) A volcano plot illustrating differentially expressed genes (DEGs) in NPCs treated with the FGFR-agonist compared to untreated NPCs. Upregulated genes are represented by red dots, downregulated genes by green dots, while non-significant genes are represented by gray dots. (B) This panel shows a functional enrichment analysis of upregulated DEGs in FGFR-agonist-treated NPCs versus untreated NPCs, utilizing gene ontology (GO) terms. It displays the top 10 significantly enriched GO terms, with the corresponding number of genes in parentheses. (C) A KEGG pathway enrichment analysis of the upregulated DEGs in FGFR-agonist-treated NPCs compared to untreated NPCs, showcasing the top 20 significantly enriched pathways, denoted by the -log10 (adjusted p-value). (D) A similar KEGG pathway enrichment analysis for downregulated DEGs in FGFR-agonist-treated NPCs compared to untreated NPCs, also showing the top 20 significantly enriched pathways. (E) This panel presents a pathway enrichment analysis of differentially expressed transcripts in FGFR-agonist-treated NPCs versus untreated NPCs, based on KEGG pathway classification, highlighting the top 20 significantly enriched pathways. (F) A KEGG pathway enrichment cnet plot for upregulated DEGs in FGFR-agonist-treated NPCs compared to untreated NPCs. The nodes symbolize KEGG pathways, and the edges signify overlapping genes between pathways. Node size corresponds to the number of genes in the pathway, while node color represents the -log10 (adjusted p-value) of enrichment.

We further analyzed the pathway enrichment analysis of differentially expressed transcripts. The results revealed that the most important signaling pathways for NPCs were upregulated, such as the Ras signaling pathway, Wnt signaling pathway, PI3K-AKT pathway, and cell cycle (**Fig. 4E**). The KEGG pathway enrichment plot analysis of upregulated genes identified three signaling networks, including cell-substrate adhesion, ERK1/2 cascade, and response to stimulus (**Fig. 4F**). Finally, we compared the transcription profiles of the NPCs treated with FGFR-agonist and bFGF and the results showed that both upregulated and downregulated gene profiles compared with the untreated NPCs were highly resembled in both conditions (**Fig. S9**). The similarity parameter of the transcription profiles between the FGFR-agonist and bFGF-treated NPCs was 0.996 (**Fig. S10**). These findings indicate that the FGFR-agonist can mimic the function of bFGF to maintain the stemness of NPCs. In conclusion, our results suggest that the FGFR-agonist can promote stem cell maintenance, regulate cell responses to growth factor stimuli, and suppress neuronal differentiation, which would further support the potential of the FGFR-agonist as an effective modulator of NPCs as bFGF.

### FGFR-agonist maintains NPCs for long-term propagation

Next, we asked if FGFR-agonist was applicable for long-term maintenance of NPCs. In our hands, the NPCs could be successfully passaged and maintained for over 50 passages using FGFR-agonist containing culturing media. The long-term propagation did not cause apparent changes in morphology or growth characteristics. We characterized the activity of FGFR-agonist in maintaining the self-renewal function of NPCs. The absence of bFGF or FGFR-agonist led to a decrease in cell survival over time, highlighting the role of FGFR-agonist for NPCs survival and self-renewal (**Fig. 5A**). FGFR-agonist effectively promoted the proliferation of NPCs in the 72-hour durations, that was comparable to the bFGF-treated condition. In contrast, the Ctrl oligo was unable to promote cell proliferation and the cell survival was greatly inhibited. These results confirmed the function of FGFR-agonist to facilitate the cell survival and self-renewal of NPCs via FGFR signaling.

**Fig. 5:**
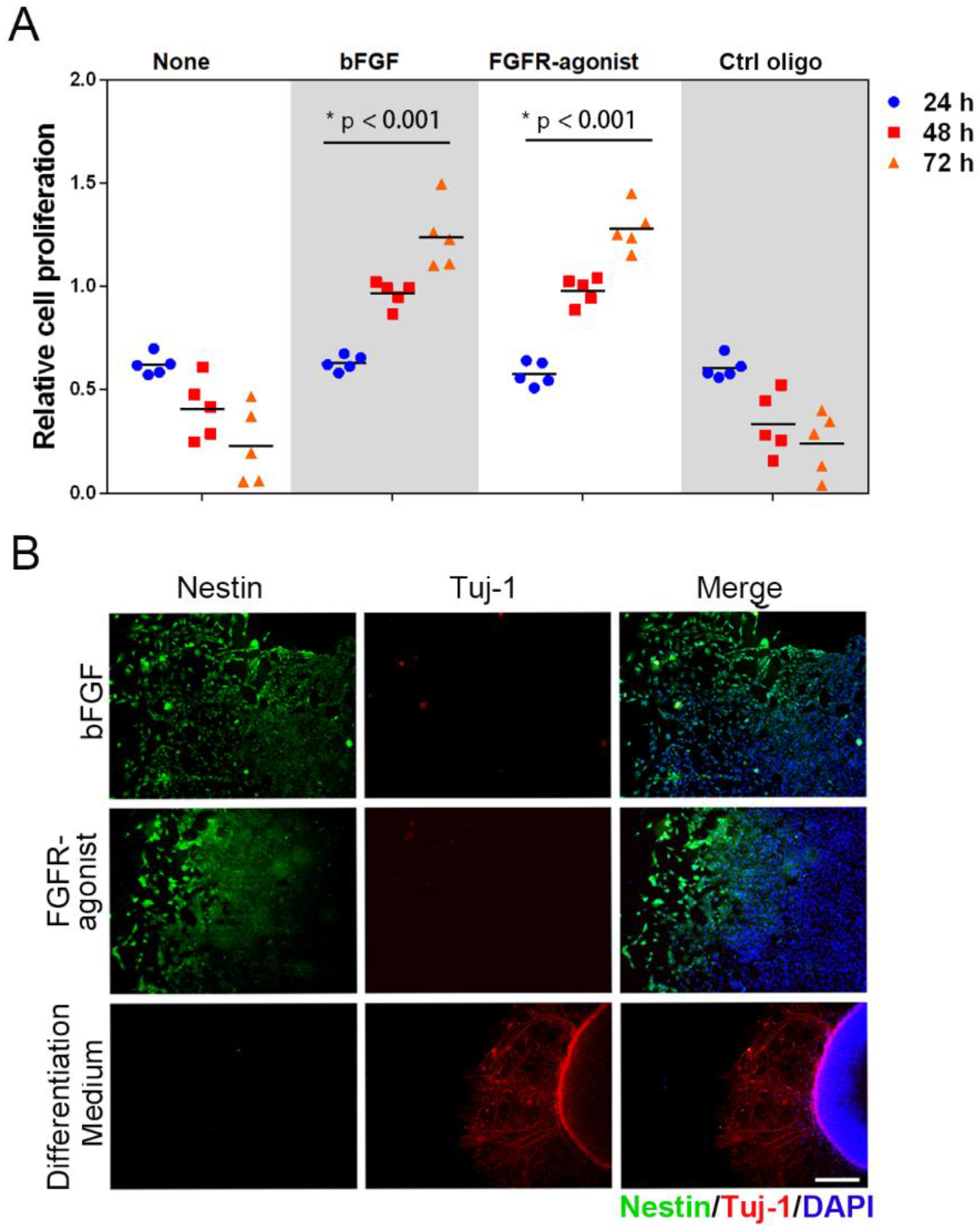
FGFR-Agonist Facilitates Sustained Propagation of NPCs with Potency for Neurogenesis. (A) This panel illustrates the influence of the FGFR-agonist on NPC proliferation. NPCs were treated with either bFGF, the FGFR-agonist, or a control oligo for a period of 72 hours, post which cell viability was assessed using the CCK8 assay. Data are depicted as mean ± SD (n = 3), with *p< 0.001 (based on a one-way ANOVA with Tukey’s multiple comparison test). (B) Immunofluorescence staining displaying Nestin (represented by green fluorescence) and Tuj-1 (indicated by red fluorescence) in NPCs subjected to treatment with bFGF, FGFR-agonist, or under a neuronal differentiation condition for 10 days. The scale bar stands for 100 μm.

Next, we explored the potency of FGFR-agonist in maintaining NPCs for neuronal differentiation. We changed the media of the FGFR-agonist-maintained NPCs with different media containing bFGF (20 ng/mL), FGFR-agonist (40 nM) or the differentiation condition, respectively. After 10 days incubation without bFGF or FGFR-agonist, the cells showed elongated morphology, resembling the neurons and immunofluorescence showed that the Tuj-1-positive signals (Red fluorescence) were strongly increased and the Nestin signal (Green fluorescence) were decreased in the neuronal differentiation group compared with the bFGF or FGFR-agonist treated groups (**Fig. 5B**). This data indicates that the NPCs cells undergoes neuronal differentiation when the FGFR-agonist absent. In contrast, FGFR-agonist treated NPCs maintained strong staining signal of Nestin, however, hardly detectable Tuj-1 signal. Together, our data validates that FGFR-agonist has similar functionality of bFGF in maintaining the multipotency of NPCs for neurogenesis.

In summary, our results demonstrate that the use of DNA-based FGFR-agonist function as an alternative to bFGF and can effectively maintain the stemness of NPCs with decoupled neuronal differentiation. Our findings highlight the potential of artificial agonists as a valuable tool for large-scale acquisition of neural cells for functional assessment and disease modeling, as well as their potential application in regenerative medicine.

## Discussion

The development of alternative culture systems for NPCs has garnered significant attention in recent years. Various strategies have been explored to enhance the scalability and consistency of NPC culture. One approach involves the use of defined media formulations devoid of serum and protein growth factors. Instead, these chemically defined media contain a precise combination of essential nutrients, vitamins, hormones, and signaling molecules to support NPC growth and maintenance [37, 38]. These chemically defined media provide better control over the culture conditions, reducing batch-to-batch variability and ensuring reproducibility of experimental results[39] . Furthermore, chemically defined media eliminate the risk of introducing immunogenic factors from serum, addressing concerns related to the safety and compatibility of NPCs for potential clinical applications [40]. Another approach to overcome the challenges in NPC culture is the use of small molecules and synthetic substitutes to replace protein growth factors. Small molecules with specific signaling properties can activate or inhibit key signaling pathways involved in NPC self-renewal, proliferation, and differentiation [41]. These molecules offer several advantages over protein growth factors, including increased stability, lower production costs, and improved reproducibility[42]. For example, small molecule agonists of Wnt signaling, such as CHIR99021, have been shown to enhance NPC proliferation and maintain their undifferentiated state[43]. Similarly, small molecule inhibitors of transforming growth factor-beta (TGF-β) signaling, such as SB431542, can promote NPC differentiation into specific neural cell lineages [44]. In addition to small molecules, synthetic substitutes for protein growth factors have emerged as a promising alternative. These substitutes can mimic the functions of native growth factors and activate their corresponding signaling pathways without the need for costly protein production [45]. For instance, DNA-based aptamers, which are short single-stranded DNA molecules, can be engineered to specifically bind and activate cell surface receptors, mimicking the effects of protein ligands[46] .These aptamers offer several advantages, including high specificity, stability, and ease of synthesis [47]. By designing aptamers that target key receptors involved in NPC self-renewal and differentiation, researchers can develop innovative strategies to modulate NPC behavior and culture them more efficiently.

We investigated the potential of a DNA-based modulator as an alternative to protein growth factors for the maintenance and expansion of human-derived NPCs. We focused on the fibroblast growth factor (FGF) signaling pathway, which plays a crucial role in NPC selfrenewal and differentiation[29] . By designing a DNA-based agonist specific to the FGFR1, we aimed to activate the downstream signaling cascade and promote NPC stemness while eliminating the limitations associated with bFGF [48]. We conducted a comprehensive evaluation of the agonist-mediated FGFR signaling and transcriptome analysis to assess the impact of the DNA-based modulator on the maintenance of NPC self-renewal over long-term culture. The successful implementation of our DNA-based modulator holds immense potential for disease modeling, drug screening, and regenerative medicine applications. By providing a consistent and scalable method for culturing NPCs, our approach opens up new avenues for studying the pathology of neurological disorders, identifying therapeutic targets, and screening potential drug candidates [49]. Additionally, the ability to generate large quantities of high-quality NPCs can facilitate the development of cell-based therapies aimed at replacing or repairing damaged neural tissue[50] .

Moreover, oligonucleotide-based therapeutics can be produced at a lower cost, making them more accessible and scalable. The use of DNA-based modulators offers several advantages over traditional protein growth factors. These modulators can be easily synthesized, modified, and optimized for specific receptor interactions[30]. They also provide greater stability and reproducibility, reducing batch-to-batch variability and experimental inconsistencies [51]. Our preliminary work seek to shed light on the advantages and feasibility of using nucleic acid agonists as an alternative to bFGF, providing valuable insights for the development of future culture systems and their practical applications in various areas of neuroscience and regenerative medicine[52]. Our findings have the potential to offer a more consistent, scalable, and cost-effective method for the large-scale production of NPCs. By addressing the limitations of current culture systems, our approach aims to provide a more consistent, scalable, and cost-effective method for large-scale NPC culture. The successful implementation of our DNA-based modulator holds immense potential for disease modeling, drug screening, and regenerative medicine applications, paving the way for future advancements in the field of neuroscience and therapeutic development.

## Supporting Information

Methods and Materials, supplementary Tables and figures.

## Acknowledgement

We would like to acknowledge the funding support from the Joint Institute of Tobacco and Health (Grant No. 2022539200340039) and the Natural Science Foundation of Hunan Province (Grant No. 2021JJ30093). We also extend our sincere gratitude to Dr. Cai Na and Mr. Ma Long from CellWay Bio Company for providing human embryonic stem cells and guidance and support in the culture and directed differentiation of NPCs.

## Corresponding Authors

## Notes

The authors declare no competing financial interest.

## Author Contributions

All authors have given approval to the final version of the manuscript.

