## Supporting Information for "Development of Synthetic Modulator Enabling Long-Term Propagation and Neurogenesis of Human-Derived Neural Progenitor Cells"

### Materials and Methods

#### hESC culture

The human embryonic stem cell line H9 (HESCs) obtained from Cellway Bio was maintained in standard hESC/hESCs medium. The medium consisted of DMEM:F12 supplemented with 20% KnockOut Serum Replacement (KOSR) and 20 ng/ml basic fibroblast growth factor (bFGF) from Invitrogen. Additional components included 1X nonessential amino acids and 110 μM 2-Mercaptoethanol. The HESCs were cultured on Matrigel-coated plates using mTeSR1 medium from Stem Cells Technologies at a temperature of 37℃. The culture plates were passaged every 6-8 days, and fresh mTeSR medium was replaced daily to maintain the cells. On the 5th day after passaging the HESCs, the cells were rinsed with sterile PBS, and a small amount of embryoid body differentiation medium was added. The HESCs colonies were gently scraped using a sterile cell scraper to detach them from the culture plate and suspend them in the medium. The suspended HESCs colonies were then transferred to cell culture dishes and cultured in the differentiation medium at 37℃ for at least 5 days. During this period, fresh medium was replaced every 2 days to support the differentiation process. To preserve the HESCs, they were stored in a cell freezing medium provided by New Cell&Molecular Biotech Co., Ltd., ensuring their long-term viability and use in future experiments.

#### Preparation of neural progenitor cells (NPCs)

After 10 days of suspension culture, the human embryonic stem cells (hESCs) formed uniform embryoid bodies with similar structures. The differentiation medium was then replaced with a mixture of N2, B27, and serum-free DMEM/F12 supplemented with Dorsomorphin and SB431542. Subsequently, the embryoid bodies were transferred to Matrigel-coated dishes and cultured at 37°C for 10 days. Throughout this period, fresh N2/B27 medium supplemented with 10 μg/L bFGF replacement was provided every 2 days. After 6 to 10 days, the cells located in the center of the embryoid bodies underwent differentiation, acquiring rosette-like structures characteristic of neural progenitor cells (NPCs). To collect the NPCs, a sterile needle was carefully used to aspirate the cells, which were then transferred to a sterile centrifuge tube. To facilitate cell pelleting, DMEM/F12 serum-free medium equivalent to 2/3 of the tube volume was added, and the cells were centrifuged at 1000 rpm. The resulting cell pellet was resuspended in a mixture of N2, B27, and serum-free DMEM/F12 supplemented with 10 μg/L bFGF. The NPCs were harvested and maintained in NPC culture medium, which consisted of either protein-based bFGF or DNA-based FGFR-agonists. For the subsequent stage of mature neuronal differentiation, N2/B27 medium without bFGF was utilized, and the NPCs were cultured for an additional 10 to 14 days. This culture condition supported the NPCs in differentiating into mature neurons. To assess stemness and neuronal differentiation, immunofluorescence staining was performed. The cells were fixed and stained with antibodies against MAP2 (Abcam) to identify neurons, as well as antibodies against Nestin (Abcam) and Pax 6 (Abcam) to identify neural progenitors. Confocal laser scanning microscopy (CLSM) (Nikon, Eclipse TE2000-E, Japan) was used to acquire images of the stained cells.

**Cell binding affinity analysis**

Cell suspensions were prepared by harvesting the cells in cell dissociation buffer. Following this, 1×$10^5$ cells were incubated with varying concentrations of the FGFR-binder or FGFR-agonist in 100 µL of buffer at room temperature. Subsequently, the cells were washed twice with 200 µL of PBS at room temperature. Finally, the cells were suspended in 200 µL of buffer and subjected to flow cytometry analysis using a flow cytometer (BD Accuri TM C6 Plus, USA). The flow cytometry data allowed for the characterization and quantification of the cells binding activity.

**FGFR activation and ERK signaling**

The oligonucleotides used in the study were provided by CellWay Bio and were purified using high-performance liquid chromatography (HPLC) to ensure their quality. To analyze FGFR phosphorylation, NIH3T3 cells and ATDC5 cells were serum-starved and stimulated with DNA-based FGFR-agonist or bFGF. The lysates were subjected to an enzyme-linked immunosorbent assay (ELISA) using a phospho-FGFR1 (Tyr653/654) kit from R&D Systems, following the manufacturer's instructions. The activation of the FGFR phosphorylation (Tyr653/654) and its downstream ERK signaling pathway was assessed by Western blot analysis, employing antibodies specific to phospho-FGFR1 (Tyr653/654), phospho-ERK1/2 (Thr202/Tyr204), and total ERK1/2 from Cell Signaling Technology. The Western blot analysis allowed for the detection and quantification of the phosphorylation levels of FGFR1 and ERK1/2, providing insights into the activation status of the signaling pathway.

**Proliferation assay**

Cell proliferation in the presence of FGFR-agonist or bFGF was determined using a Cell Counting Kit-8 (CCK-8) assay obtained from New Cell&Molecular Biotech Co., Ltd. Neural progenitor cells (NPCs) were seeded in a 96-well plate at a density of 1x$10^5$ cells per well and incubated for 24, 48, and 72 hours. Following the respective incubation periods, the CCK-8 reagent was added to each well, and the absorbance at 450 nm was measured using a microplate reader. The CCK-8 assay provides a quantitative assessment of cell viability and proliferation by measuring the metabolic activity of the cells, with higher absorbance values indicating increased cell proliferation.

**Wound scratch assay**

Cells (1x$10^5$) were seeded in a 12-well plate at a density that allowed them to reach approximately 70-80% confluence as a monolayer after 24 hours of incubation. Subsequently, wounds were created in each cell monolayer by carefully scratching them using a sterile 200 µL pipette tip. After scratching, the wells were gently washed twice with medium to remove any detached cells. The cells were then incubated with either bFGF or FGFR-agonist. The rate of wound healing was monitored and imaged at 24 and 48 hours using a Cytation 5 microplate reader equipped with a positioning system. The images captured at different time points were analyzed to measure the extent of wound closure, providing quantitative data on the rate of cell migration and wound healing.

### Transcriptomic analysis

Total RNA was extracted from the neural progenitor cells (NPCs) using TRIzol reagent (Invitrogen) following the manufacturer's instructions. The extracted RNA samples were then subjected to RNA sequencing using a nanopore sequencing platform (Oxford Nanopore Technologies). The generated sequencing data were processed and analyzed using a customized bioinformatics pipeline. To gain insights into the functional significance of the differentially expressed genes (DEGs), gene ontology (GO) analysis and Kyoto Encyclopedia of Genes and Genomes (KEGG) pathway analysis were performed. GO analysis provided information on the biological processes, cellular components, and molecular functions associated with the DEGs. KEGG pathway analysis identified the signaling pathways and networks in which the DEGs were significantly enriched. These analyses aimed to identify genes and pathways involved in stem cell maintenance and neural differentiation. By comparing the expression profiles of NPCs cultured in the presence of artificial agonists or bFGF, we aimed to elucidate the similarities and differences in gene expression patterns and pathway activation. The results of the bioinformatics analysis provided valuable insights into the molecular mechanisms underlying the effects of the artificial agonists and bFGF on stem cell maintenance and neural differentiation.

### Long-term NPCs maintenance and neuronal differentiation

To assess the differentiation potential of neural progenitor cells (NPCs) cultured in the presence of the FGFR-agonist or bFGF, the cells were passaged and maintained for over 50 passages. Subsequently, the cells were subjected to a differentiation assay by withdrawing bFGF or the FGFR-agonist and replacing it with a neuronal differentiation medium. After 10 days of differentiation, the cells were fixed and immunostained with specific antibodies against neuronal marker Tuj-1 (Abcam) and NPC marker Nestin (Abcam). Fluorescence microscopy analysis was performed to visualize and evaluate the presence of differentiated neurons (Tuj-1-positive cells) and the maintenance of neural progenitor cell characteristics (Nestin-positive cells) in the cultures. Confocal laser scanning microscopy (CLSM) (Nikon, Eclipse TE2000-E, Japan) was used to acquire images of the stained cells. The immunostaining results provided insights into the effect of the FGFR-agonist compared to bFGF in promoting neuronal differentiation and maintaining the NPC phenotype.

### Statistical analysis

All experiments were conducted with triplicate samples, and the data are presented as the mean ± standard deviation (SD). Statistical analyses were performed using unpaired two-tailed Student's t-test or one-way analysis of variance (ANOVA), followed by Tukey's post hoc test, as appropriate. A p-value of less than 0.001 was considered statistically significant. Statistical significance is indicated as follows: *$p < 0.001$, **$p < 0.0001$. If the p-value was greater than 0.05, it is indicated as "n.s." (no significance).

**Table S1**

| Name | Sequence (5'---3') |
|---|---|
| FGFR-binder | GCC GCG TCT TTA TGG CTG GGG ATG GTG TGG GTT GCG GC |
| FGFR-agonist | GCC GCG TCT TTA TGG CTG GGG ATG GTG TGG GTT GCG GCG CCG CGT CTT TAT GGC TGG GGA TGG TGT GGG TTG CGG C |
| Ctrl oligo | CGG CGT TGG GTG TGG TAG GGG TCG GTA TTT CTG CGC CGC GGC GTT GGG TGT GGT AGG GGT CGG TAT TTC TGC GCC G |

**Table S2**

| Samples | DEG | Upregulated | Downregulated |
|---|---|---|---|
| None vs bFGF | 189 | 75 | 114 |
| None vs FGFR-agonist | 208 | 111 | 97 |

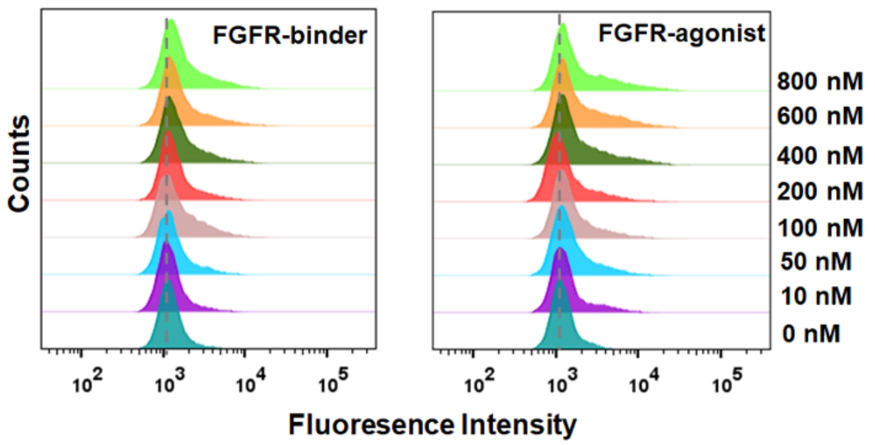


**Fig. S1: Examination of FGFR-Binder and FGFR-Agonist Binding Affinity with NIH3T3 Cells via Flow Cytometry.**

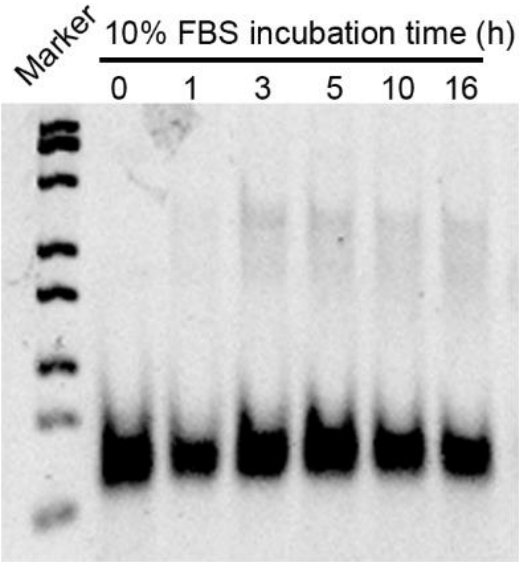


**Fig. S2: Serum Stability Assessment of FGFR-Agonist.** Different DNA oligonucleotides were incubated with 10% fetal bovine serum (FBS) for varying durations, followed by PAGE gel electrophoresis to evaluate stability.

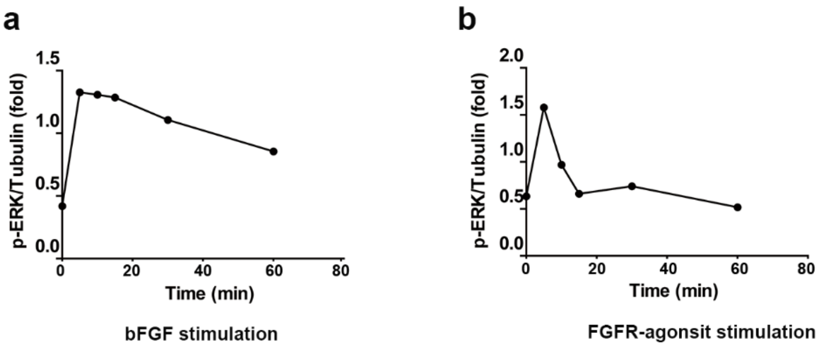


**Fig. S3. Time-Dependent ERK Phosphorylation Induced by bFGF (a) or FGFR-Agonist(b).** Serum-starved ATDC5 cells were stimulated with either bFGF (20 ng/mL) or FGFR-agonist (40 nM) for specified time intervals (5, 10, 15, 30, 60 minutes). Phosphorylation levels of ERK1/2 at Thr202 and Tyr204 were assessed via western blotting. Quantitative analysis was performed using Image J software to represent the kinetics of ERK1/2 phosphorylation in response to different stimulations.

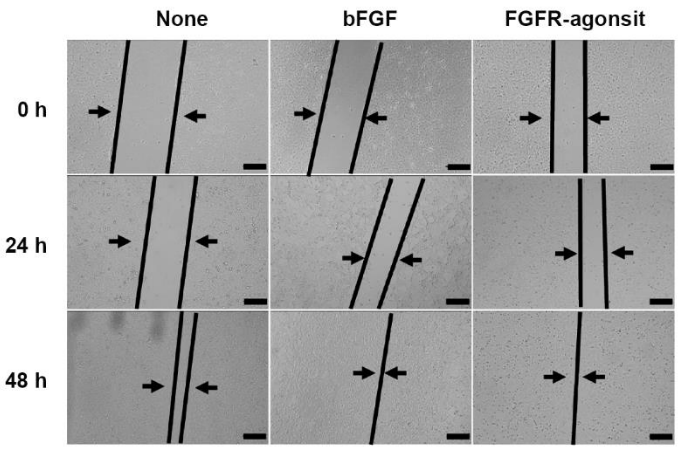


**Fig. S4：ATDC5 Cell Wound-Closure Events Monitoring.** Cells were treated with bFGF, FGFR-agonist, or FGFR-binder for 2 days, and wound-closure events were tracked under light microscopy with images taken at 0 h, 24 h, and 48 h. Black lines indicate boundaries between scratched monolayers. Scale bar: 200 μm.

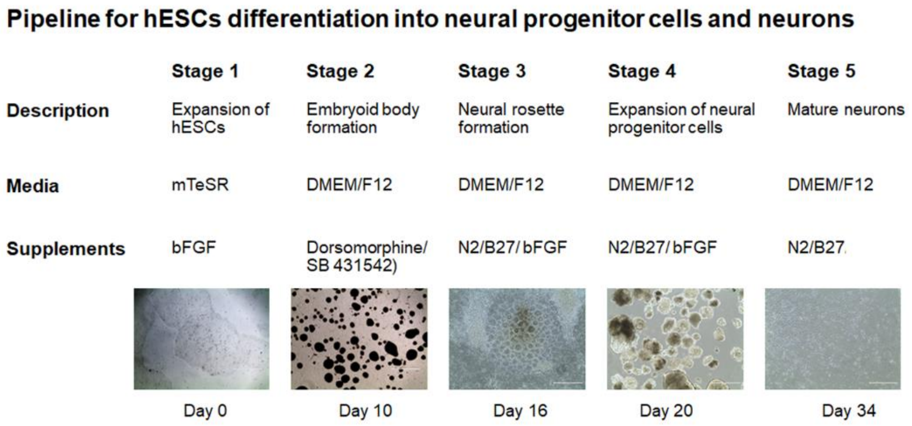


**Fig. S5: Schematic Diagram of the Experimental Procedure for Neuronal Differentiation from hESCs.** The necessary supplements and media for each stage are listed. Representative images of hESCs (Stage 1), Embryoid Body (Stage 2), Neural Rosette (Stage 3), Neural Progenitor Cells (Stage 4), and Mature Neurons (Stage 5) were captured under light microscopy at indicated time points. Scale bar: 500 μm.

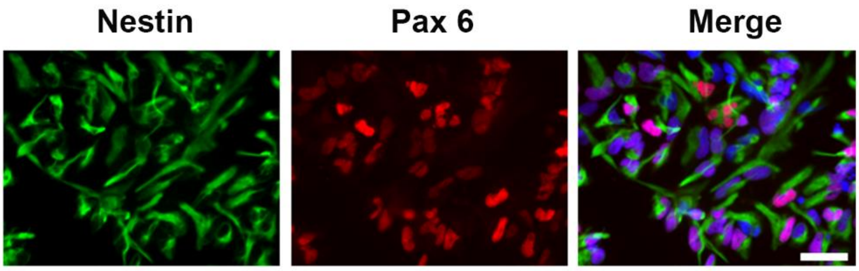


**Fig. S6: Immunofluorescence Staining of hESC-Derived NPCs.** Cells were fixed and stained with antibodies against Nestin (Green) and Pax 6 (Red). Nuclei were counterstained with DAPI (Blue). Scale bar: 50 μm.

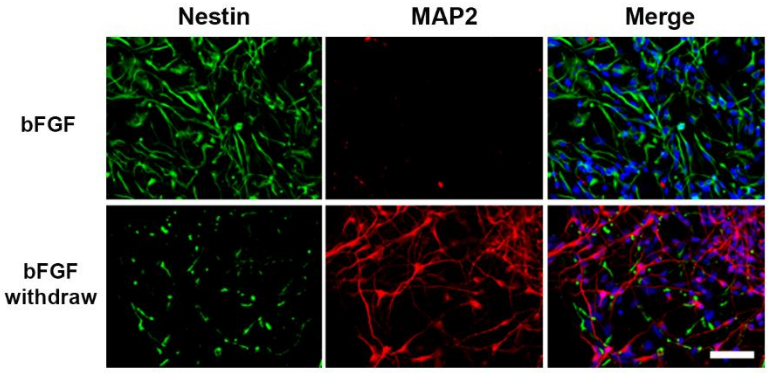


**Fig. S7**. **Immunofluorescence Staining of NPCs Following 10-day Treatment with or without bFGF.** NPCs were fixed and stained for Nestin (Green) and MAP2 (Red). Nuclei were counterstained with DAPI (Blue). Scale bar: 50 μm.

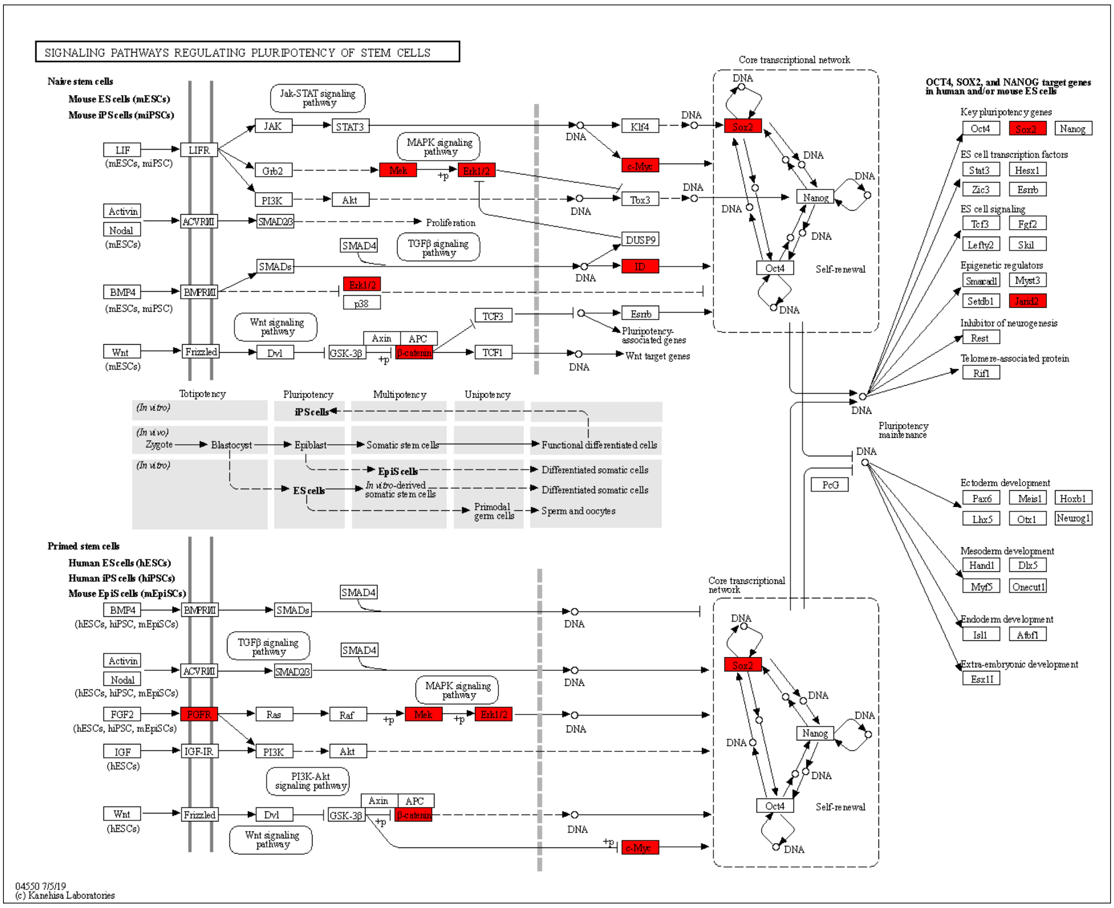


**Fig. S8: KEGG Pathway Analysis Illustrating Enhanced Stemness Markers in FGFR-Agonist Treated Group.** The KEGG pathway analysis (ko04550) is employed to pinpoint several key stemness marker genes that display elevated expression following FGFR-agonist treatment. The key upregulated genes including FGFR, MERK, ERK1/2, Jarid2, SOX2 and c-Myc, that were highlighted with red in the KEGG map.

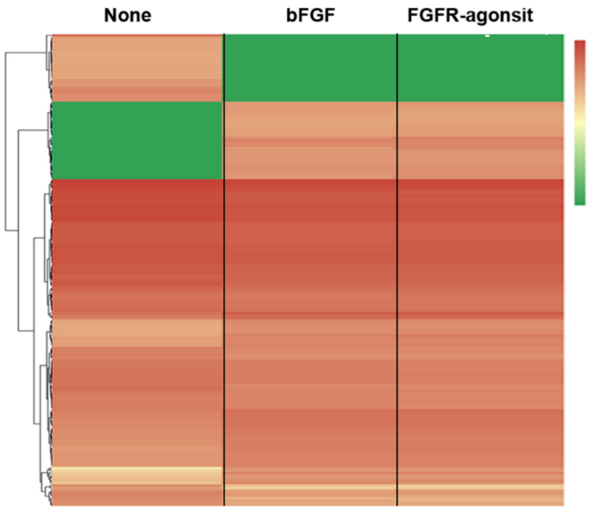


**Fig. S9. Heatmaps of Gene Profiles in NPCs Treated with bFGF or FGFR-Agonist.**

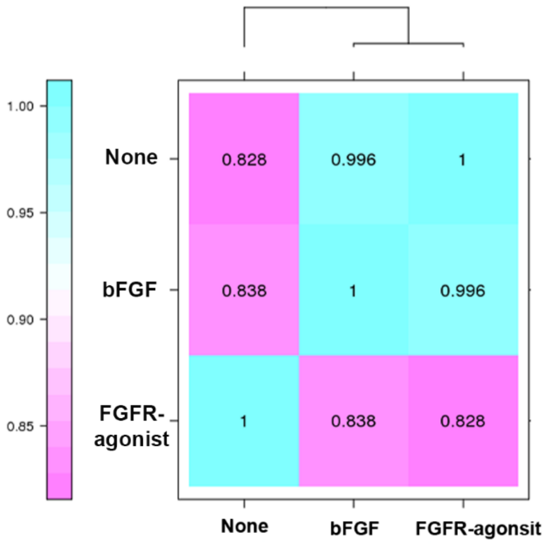


**Fig. S10. Correlation Heatmap of Gene Expression Levels Among Samples, Including None, bFGF, and FGFR-Agonist.**
